# Unsupervised explainable AI for simultaneous molecular evolutionary study of forty thousand SARS-CoV-2 genomes

**DOI:** 10.1101/2020.10.11.335406

**Authors:** Toshimichi Ikemura, Kennosuke Wada, Yoshiko Wada, Yuki Iwasaki, Takashi Abe

## Abstract

Unsupervised AI (artificial intelligence) can obtain novel knowledge from big data without particular models or prior knowledge and is highly desirable for unveiling hidden features in big data. SARS-CoV-2 poses a serious threat to public health and one important issue in characterizing this fast-evolving virus is to elucidate various aspects of their genome sequence changes. We previously established unsupervised AI, a BLSOM (batch-learning SOM), which can analyze five million genomic sequences simultaneously. The present study applied the BLSOM to the oligonucleotide compositions of forty thousand SARS-CoV-2 genomes. While only the oligonucleotide composition was given, the obtained clusters of genomes corresponded primarily to known main clades and internal divisions in the main clades. Since the BLSOM is explainable AI, it reveals which features of the oligonucleotide composition are responsible for clade clustering. The BLSOM has powerful image display capabilities and enables efficient knowledge discovery about viral evolutionary processes.

## Introduction

To confront the global threat of COVID-19^1,2^, many SARS-CoV-2 genome sequences have been rapidly decoded and promptly released through the GISAID database^3^. To characterize this virus in various ways, we must implement diverse research methods such as AI (artificial intelligence), that are suitable for big data analyses. Unsupervised machine learning can obtain new information from big data without particular models or presumptions and is highly desirable for mining big data. We previously established a BLSOM (batch-learning self-organizing map) for oligonucleotide compositions, which can reveal various new characteristics of genome sequences^4,5^.

Oligonucleotide composition varies significantly among species, even those with the same genome G+C%, and is called the genome signature^6^. When we constructed a BLSOM for oligonucleotide compositions in fragment sequences (e.g., 10 kb) from a wide variety of species, the sequences were clustered (self-organized) primarily according to species, despite no species information being used during machine learning^5,7^. Importantly, the BLSOM is suitable for large-scale analysis and has been used to analyze five million genomic fragments from over one thousand genera^8^. In addition, the BLSOM is explainable AI and can reveal the drivers of species-specific clustering (self-organization).

Many host factors (e.g., nucleotide pools, proteins and RNAs) and antiviral mechanisms (e.g., antibodies, cytotoxic T cells and interferons) are involved in viral growth and infection^9,10^. Since human cells may not present ideal growth conditions for zoonotic viruses that have invaded from nonhuman hosts, efficient growth and human-human transmission will likely require changes in the viral genome. To study this viral adaptation, we previously analyzed time-series changes in the mono- and oligonucleotide compositions of four zoonotic RNA viruses (influenza virus^11,12^, Zaire ebolavirus^13^, MERS coronavirus^13^ and SARS-CoV-2^14^) and identified time-series directional changes that were detectable even on a monthly basis.

In the case of fast-evolving RNA viruses, diversity within the viral population arises rapidly as the epidemic progresses and subpopulation structure forms, and the GISAID consortium has defined seven main clades, and Mercatelli and Giorgi (2020)^15^ have recently conducted a large-scale search for prevalent mutations worldwide. In the present study, over 40,000 genomes of SARS-CoV-2 are analyzed by using BLSOMs with oligonucleotides of various lengths. The BLSOM is a sequence alignment-free method, and during machine learning, only the oligonucleotide composition of each viral genome is given. Therefore, clustering is performed basing only on the similarity of oligonucleotide compositions. By trying various oligonucleotide lengths, we obtained conditions for separating the known clades with high accuracy. Since the BLSOM is explicable AI, it can identify the features of oligonucleotide compositions responsible for the separation. Since the BLSOM method is based on a completely different principle than conventional clade assignment based on sequence alignment, additional information can be obtained.

## Results

### BLSOM for 1 ∼ 6-mers

In the present study, over forty thousand SARS-CoV-2 genomes, which have been isolated from December 2019 to June 2020, were analyzed; polyA-tail was removed prior to all analyses. Figure 1 shows the BLSOM results for the mono-to hexanucleotide compositions in the viral genomes; importantly, only the composition was used in the learning process. The total number of nodes (grid points) was set to 1/20 of the total number of viral genomes (40,450); therefore, each node had an average of 20 genomes. After learning, to determine if the separation by the BLSOM was related to known clads, grid points containing genomes of a single clade were colored to indicate each clade, and grid points containing those of multiple clades were displayed in black. Figure 1 shows that most grid points in the mononucleotide BLSOM are black, indicating that the genome sequences are not separated by clade. However, for greater than dinucleotide lengths, the separation by clade gradually becomes clear. While each node has 20 genomes on average, and even though grids containing at least one genome belonging to another clade are marked in black, a major portion of the grids in the BLSOM for 4- to 6-mers are colored, showing the good classification power of the BLSOM. We thus tested oligonucleotides longer than 6-mers.

**Fig. 1.**
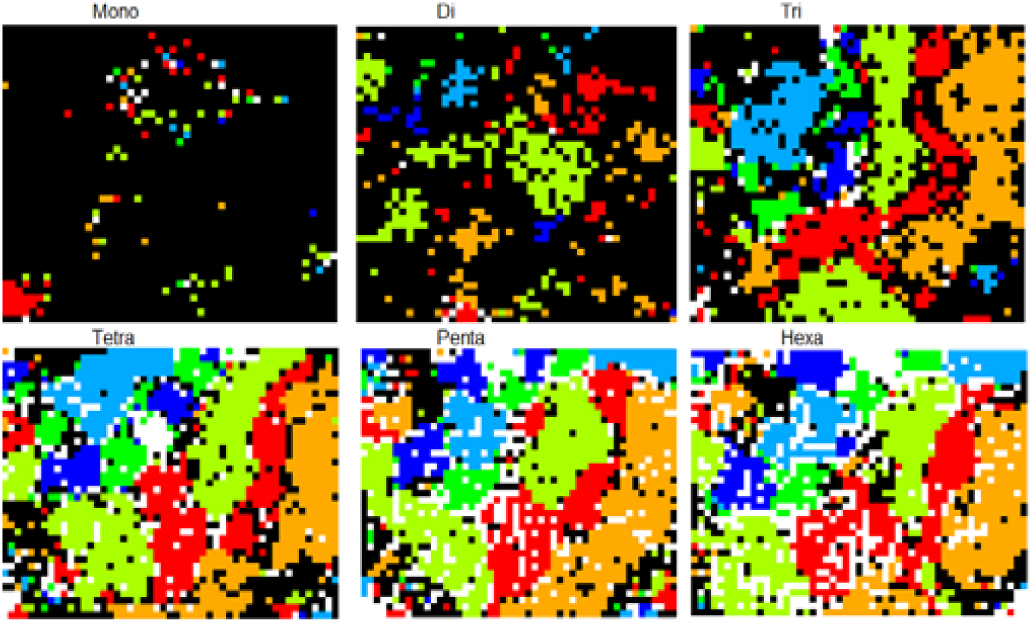
BLSOMs for 1∼6-mers. BLSOMs were constructed for mono- to hexa-nucleotide compositions in 40,450 genome sequences. The total number of nodes was set to 1/20 of the total number of viral genomes. Grid points that include sequences from more than one clade are indicated in black, and those containing sequences from a single clade are indicated in a clade-specific color: G (▪), GH (▪), GR (▪), L (▪), S (▪) and V (▪). Sequences of the O clade (other and unclassified clades) were included in the BLSOM calculation but excluded from the final display; i.e., if a sequence belonging to the O clade is mixed with sequences belonging to a main clade, the node is colored according to the main clade.

### BLSOM for 7-mers

For 7-mers, the BLSOM handles 16,384 (the 7th power of 4) variables, and for efficient analysis, some modification of PCA used to set the initial state for machine learning is required, as described in the Methods. The BLSOM with this minor modification provides good separation by clade (Fig. 2ai).

**Fig. 2.**
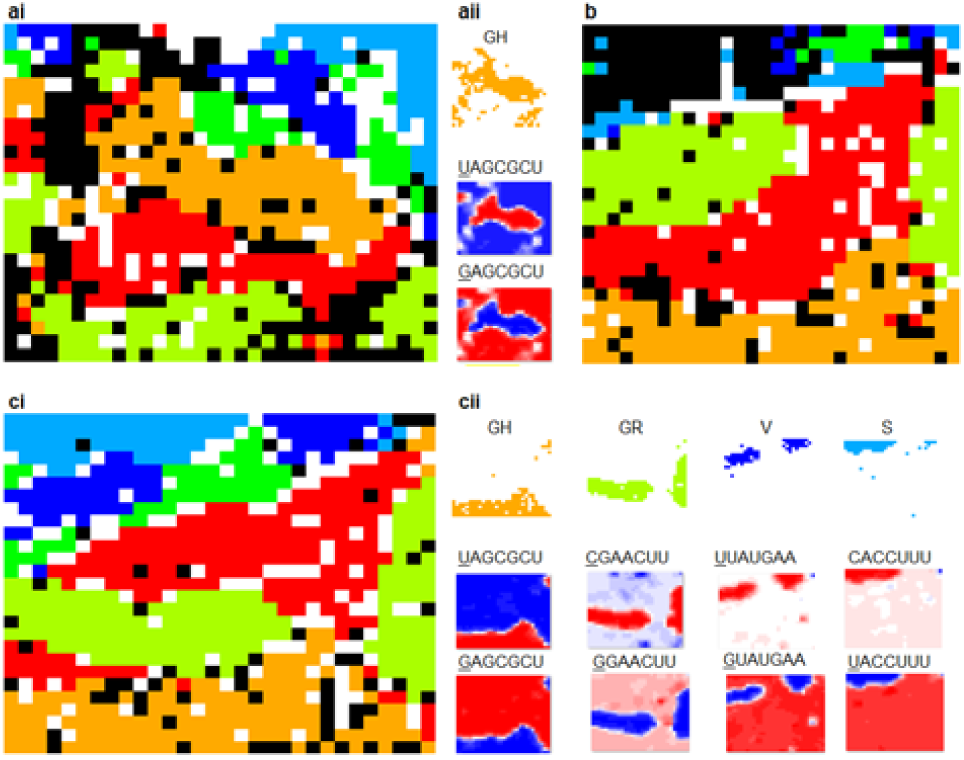
BLSOM for 7-mers. (**a**) BLSOM and heatmap. The total number of nodes was set to 1/50 of the total number of viral genomes. (i) Nodes are colored as described in Fig. 1. (ii) The top panel shows the GH territory (▪) on the BLSOM. The middle and bottom panels show heatmaps of <u>U</u>AGCGCU and <u>G</u>AGCGCU, respectively, which differ only in the underlined base. (**b**) BLSOMs for 255 different 7-mers. Nodes are colored as described in Fig. 1. (**c**) BLSOMs for 377 different 7-mers. (i) Nodes are colored as described in Fig. 1. (ii) The top four panels show the four different territories colored as described in Fig. 1. The middle and bottom panels show heatmaps of four pairs of 7-mers with a one-base difference.

BLSOM is explainable AI and can provide information about the oligonucleotides responsible for the clustering (Fig. 2aii); the representative vector for each node is composed of 16,384 variables, and the contribution levels of the variables at each node can be visualized by a heatmap: high (red), moderate (white) and low (blue)^16^. In Fig. 2aii, two examples of 7-mers are presented; <u>U</u>AGCGCU has a high occurrence (red) primarily in GH zones (mandarin orange), whereas <u>G</u>AGCGCU, which differs from the former by one underlined base, has a low occurrence (blue) there. When considering all 7-mer patterns (refer to Supplementary Fig. S1), we observed multiple cases in which occurrence levels visualized as red/blue were reversed for a pair of 7-mers with a one-base difference, as shown in Fig. 2aii. Notably, these reverse patterns were primarily observed for individual clade territories, indicating that one-base differences may be related to mutations involved in clade separation.

### Necessity for dimension reduction

Notably, most 7-mers existed in multiple copies in the viral genomes; therefore, a one-base difference giving the red/blue reverse pattern could not be connected uniquely to a mutation in the viral genome. In other words, extending the oligonucleotide length until most k-mers were present as one copy per genome should allow us to connect one-base differences to mutations. Accordingly, we calculated the occurrences of long oligonucleotides in the viral genomes, and when the length was extended to 15-mers, most showed one copy per genome. The 15-mers, however, included over one billion types (the 15th power of 4), and even for those that appeared in the viral genomes, there were more than 0.3 million types. A strategy for dimension reduction is essential for efficient analyses; then we examined the effects of dimension reduction on clade separation by testing 7-mers again.

In Supplementary Fig. S1, most 7-mers are red/blue all over and very locally dotted with blue/red, respectively. These 7-mers are thought to correspond to sequences that have not mutated in most strains or that arose by mutations but remain in only a small number of strains. The oligonucleotides whose frequency in the viral population has not significantly changed with time should not be involved in the formation of main subpopulations, such as the main clades. To exclude these numerous 7-mers and select those that have changed significantly in occurrence during the pandemic, we first tabulated viral strains for each month of collection and calculated the 7-mer occurrence frequency therein. To compare the frequencies in the June (2020) and December (2019) populations, we next selected 7-mers whose frequency in the June population increased/decreased by at least 0.1 compared with that in the December population. We constructed a BLSOM with these 255 selected 7-mers (Fig. 2b). Even using a very minor portion of the 7-mers (255/16384 = 0.016), good separation by clade was observed, but the mutual separation of L, S and V clades, which are prevalent in Asia, was poor. This should be because strains belonging to these clades were very minor in the June population.

To study evolutionary processes throughout the epidemic phase, we must consider an intermediate phase and choose the March population here; then we selected 7-mers whose frequency in the March population increased/decreased by at least 0.1 compared with that in the December population. The 334 obtained 7-mers were combined with the June 7-mers used in Fig. 2b. After excluding duplicates, we constructed a BLSOM for the 377 remaining 7-mers. Importantly, the BLSOM separation (Fig. 3ci) was clearer than that observed when using 16384 types of 7-mers (Fig. 3ai), and the red/blue reverse pattern was clearly connected to clade separation; for four clades, Fig. 2cii presents a pair of red/blue patterns corresponding to a pair of 7-mers with a one-base difference, and Supplementary Fig. S2 presents all red/blue patterns. Since the dimension-reduction strategy appeared to be useful, we applied it to the 15-mer BLSOM.

**Fig. 3.**
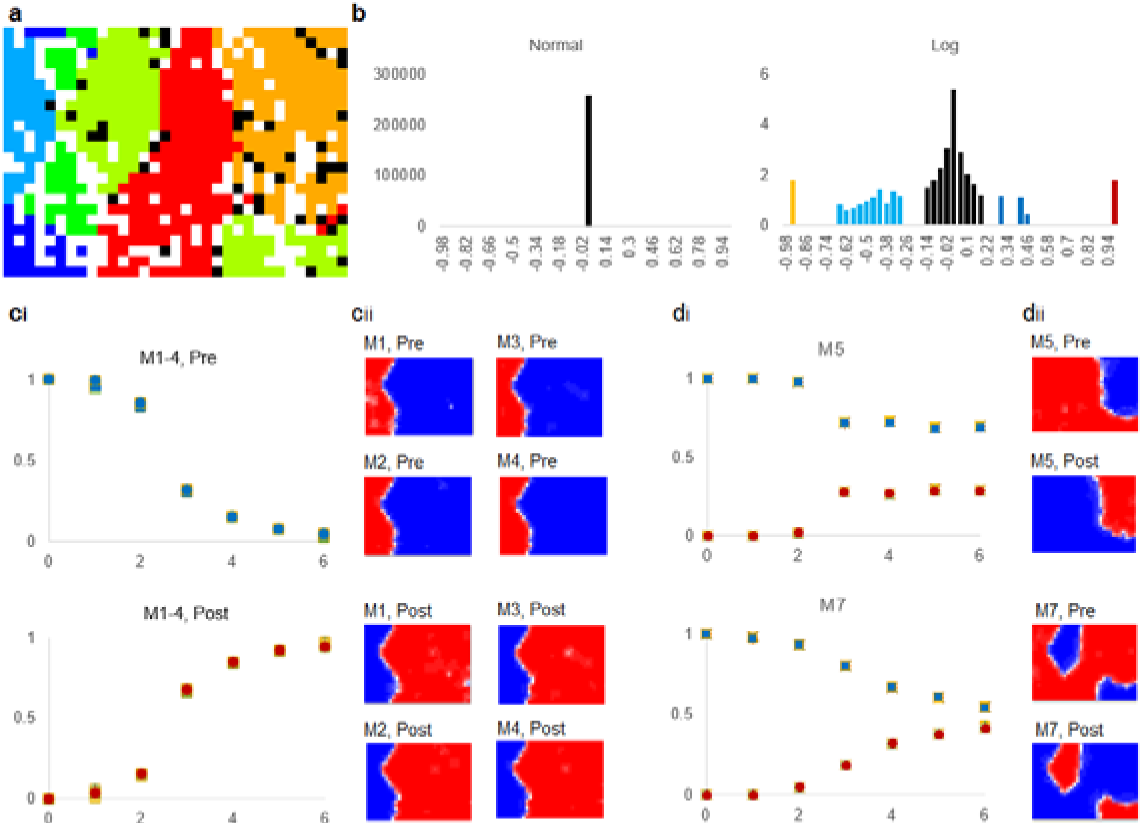
BLSOM and histogram analysis of 15-mers. (**a**) BLSOM of 15-mers. The number of nodes was set as described in Fig. 2a. (**b**) Histogram of the increase/decrease level of each 15-mer frequency in the June population compared to the December population. The vertical axis shows normal numbers or logarithms (Log). Here, nonexistence in the logarithmic display is shown expediently as 0; the dark- and light-brown (blue) bars in the histogram indicate data showing the largest (the second largest) class of the increase/decrease, respectively. (**c**) Changes in the monthly occurrences of 15-mers and their heatmap patterns. (i) The upper and lower panels plot the monthly occurrences of 15-mers according to the elapsed month, which are related to the M1∼4 mutations in Table 1 (see also Supplementary Fig. S3) and indicated by dark and light brown in b, respectively. Since there is little difference among the sixty 15-mers, the relationship between each 15-mer and the colored symbol was not described. (ii) Heatmaps of four pairs of 15-mers with a one-base difference; the four in the upper and lower panels show heatmaps of the pre- and postmutation sequences. (**d**) (i) The upper and lower panels plot monthly occurrences of a group of fifteen 15-mers belonging to the second largest class specified by dark and light blue, respectively. Here, the pre- and postmutation 15-mers are plotted in the same figure: M5 and M7 mutations in Table 1. (ii) Heatmaps of two pairs of 15-mers with one-base differences are placed next to the corresponding time-series diagram.

### BLSOM for 15-mers

Most 15-mers have only one copy in the viral genome as mentioned above. More precisely, ten 15-mers have had two copies since December 2019, and the two copies are present in almost all strains until June, so these 15-mers are unrelated to mutations with significant changes in population frequency during the pandemic. As performed for 7-mers in Fig. 2c, we selected 15-mers whose frequency in the March/June population increased/decreased by at least 0.1 compared with that in the December population and obtained 587 different 15-mers.

**Table 1.**
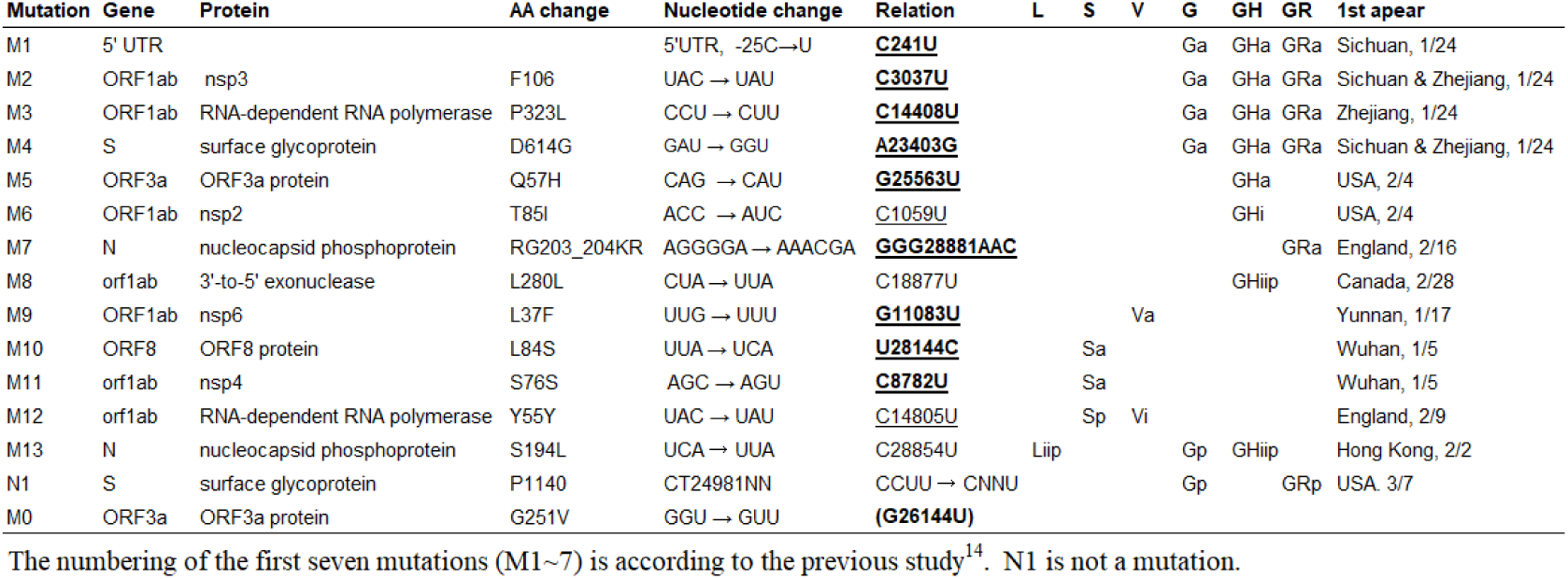

Because a maximum of one copy of these 15-mers was present in the genome, the occurrence frequency of a given 15-mer in a certain population corresponds to the frequency of the strains having the 15-mer sequence. The BLSOM with the 587 different 15-mers (Fig. 3a) showed good separation by clade. While each node has 50 genomes on average, and even though grids containing at least one genome belonging to another clade are marked in black, a major portion of the grids are colored. Next, we will examine why this good separation was obtained by focusing only on these 15-mers.

### 15-mers with rapidly changing population frequencies

The novel characteristic of evolutionary study of this fast-evolving virus is near-future prediction and verification on a monthly basis. We previously performed a time-series analysis of 20-mer occurrences in over ten thousand SARS-CoV-2 strains isolated from December 2019 to April 2020 and identified seven mutations rapidly increasing in population frequency^14^; for all 20-mers, we created a histogram of increase levels in the April population compared with the December population and identified the rapidly increasing mutations. Here, we conducted a similar histogram analysis of over 250,000 types of 15-mers found in over forty thousand viral genomes; we analyzed the occurrence level of each 15-mer on a monthly basis. Figure 3b is a histogram of the differences in 15-mer frequency between the June and December populations.

On the horizontal axis, the frequency difference relative to December is displayed, and on the vertical axis, the number of 15-mer types with the frequency difference in a 0.04 range is displayed. In the first histogram, the numbers of 15-mer types are displayed as normal values. In the center, where the frequency difference is close to zero, there is a very high peak, showing that most 15-mers underwent little change in population frequency from December; i.e., most 15-mer sequences did not mutated, or if they did, they did not spread significantly in the population. These 15-mers are not responsible for formation of main subpopulations such as main clades.

In the other histogram, the vertical axis is displayed logarithmically to show low values. Interestingly, there are clear characteristic peaks near both ends where the increase/decrease in frequency exceeds 0.9 (colored in dark and light brown, respectively), which has a logarithmic value of 1.778 (= 60); i.e., 60 types of 15-mers that belong to the dark/light brown peak have drastically increased/decreased in population frequency, respectively. The BLASTn analysis of a total of 120 types of the 15-mers against a standard viral genome sequence^17,18^ showed that these 120 types were related to four mutations, which correspond to four of the seven mutations that were previously found to be rapidly increasing in population frequency^14^. Since the fifteen 15-mers with one mutation can be represented as one 29-mer sequence, the pre- and postmutation sequences of the 29-mers are presented in Supplementary Fig. S3.

In the upper and lower panels of Fig. 3ci, the monthly population frequencies of the increasing/decreasing 15-mers are arranged according to the elapsed month. Since these 15-mers are the pairs produced by mutations, the two panels show symmetrical time-series changes. Notably, these mutations were previously found to increase monotonically to a frequency of 0.84 until April and continued to increase to 0.94 until June, supporting the monotonic increase tend reported in the previous study^14^.

### Reverse red/blue patterns for 15-mers

In the BLSOM shown in Fig. 3a, the representative vector for each node is composed of 587 variables, and the contribution level of each variable at each node can be visualized by a red/blue heatmap (Fig. 3cii). Since fifteen pairs of 15-mers related to one mutation showed primarily the same red/blue patter, one example of each mutation is presented in Fig. 3cii; the upper and lower panels show the patterns for the pre- and postmutation 15-mers, respectively. Notably, for the four mutations, very similar red/blue reverse patterns were observed, which correspond to the separation of G (red), GH (mandarin orange) and GR (yellow green) territories from L (green), S (cyan) and V (blue) territories of the Asian-type. In addition to dark and light brown peaks in the logarithmic histogram, characteristic peaks are also observed at positions apart from the central peaks, whose increases/decreases exceed 0.2, and are specified by dark/light blue; the number of peaks on the increase side (dark blue) is smaller than that on the decrease side (light blue), and the BLASTn analysis of 15-mers belonging to the dark blue peaks revealed that they are related to the two mutations (M5 and M7 in Table 1), which were also reported previously^14^. In Fig. 3di, the time-series changes of increasing and decreasing 15-mers are plotted in the same panel.

### Mutations involved in main clade separation

We next explain details of the six mutations described in Fig. 3. Table 1 lists the nucleotide and amino-acid changes and the clade territories in the BLSOM where the mutated sequences are located (i.e., the red region in the heatmap) along with the day of first isolation of the strain with the respective mutation. When the red region corresponds primarily to the entire territory of a certain clad, the suffix “a” is added, and when the red region is a part of a certain clade territory but is a continuous zone, “i” and “ii” are used to distinguish the separate zone. In addition, when the red areas are scattered in a clade, they are noted by “p”. The sequences with the first four mutations (M1-4) are specified as Ga, GHa and GRa, because the mutated sequences exist in the entire areas of G, GH and GR; these mutations are thought to relate to the separation of G and its offspring GH and GR, which are prevalent in Europe, from the Asian-types L, S and V. The first isolation date for these mutations was 1/24 in Sichuan or Zhejiang in China, and the second isolation date was 1/28 in Germany or Lishui in China. The sequences with the M5 or M7 mutation were localized in GHa or GRa, respectively, showing that the mutations should relate to the separation of GH and GR from G; the first isolation date of M5 and M7 was 2/4 in the USA and 2/16 in England, respectively.

### Consistency in identifying mutations with the phylogenetic clustering method

With a phylogenetic method using NUCMER^19^ for sequence alignment, Mercatelli and Giorgi (2020)^15^ have conducted a large-scale search for common mutations worldwide and compared them with ten known mutations that have been associated with main clade separation by the GISAID consortium. The genomic positions of the ten mutations are shown in bold in the Relation column in Table 1, and when our identified locations matched the mutations reported by Mercatelli and Giorgi (2020)^15^, they were underlined. Our analysis relies only on BLSOM, histogram and time-series analyses of oligonucleotide composition, all of which are sequence alignment-free methods. The degree to which these different methods give similar results is important for knowing the reliability of the method. The six mutations assigned in the analysis of Fig. 3c and d were found to be among the ten known mutations and thus were underlined. Notably, in Fig. 3c and d, the 15-mers whose increase/decrease was remarkable to others and exceeded 0.2 in the histogram of Fig. 3bii were analyzed. Since the BLSOM shown in Fig. 3a targeted the 15-mers with an increase/decrease of at least 0.1, it contains all of these prominent 15-mers, resulting in a good separation according to main clades. In addition, since the BLSOM targeted the 15-mers with an increase/decrease of at least 0.1, the time-series change and heatmap pattern can be obtained for more 15-mers than those analyzed in Fig. 3c and d, and Fig. 4a∼g present the additional results showing the reverse pattern for both the time-series change and the heatmap diagram. In Figs. 4a for M6 and 4b for M8, differential parts of the GH territory are red, showing the internal branching of GH; M6 first appeared on 2/4 in the USA, and M8 appeared on 2/28 in Canada. In Fig. 4c for M9, the red area covers the V territory, and this mutation was first isolated on 1/17 in Yunnan and among the ten mutations. Figures 4d for M10 and 4e for M11 show that the red area covers the S territory, indicating that the two mutations, which first appeared in the same strain isolated on 1/5 in Wuhan, are related to the S separation. Since M11 (but not M10) is a synonymous mutation and has been lost in some S-clade strains since 1/23, M11 may be a hitchhiker-type neutral mutation that has increased with M10.

**Figure 4.**
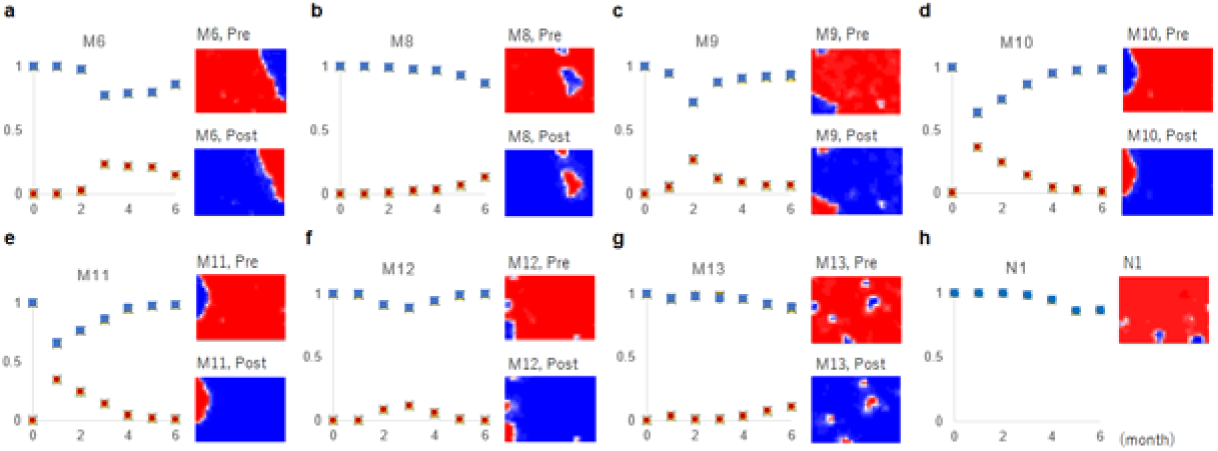
Change in monthly occurrences of 15-mers and their heatmaps. Nine time-series diagrams and heatmaps are presented as described in Fig. 3d.

In the case of M12 (Fig. 4f), a part of the V and S territories, as well as a very minor part of the GR territory, is red; this is a synonymous mutation of Tyr and may have occurred independently. In the case of M13 (Fig. 4g), it is red locally at L, G, and GH; for L, it first appeared on 1/23 in Guangdong, for G on 3/5 in the USA and for GHiip on 3/15 in India and Saudi Arabia. Notably, the strains that appeared in the GHiip territory are prevalent in Asia and overlap, in the BLSOM, with or are adjacent to the Asia-prevalent G strains that also have the M13, showing that the position within on the map reflects similarities in the presence of mutations other than those contributing to the main clade separation. Since this is a nonsynonymous mutation from Ser to Leu and continues to increase monotonically over time, it appears to be a functionally beneficial mutation. This mutation (M13) and M8, which are not among mutations reported by Mercatelli and Giorgi (2020)^15^, have increased in frequency mainly since May. Collectively, the three mutations (M7, 9 and 11) are also among the ten mutations related to the main clade separation; therefore, nine mutations analyzed so far were among the ten mutations, and the remaining one (M0 in Table 1) will be discussed in the Discussion section. Notably, AI can separate the V clade from others without the 15-mers related to M0, and this type of information should be important for identifying the mutation essential for separation of the respective clades.

### Decreasing 15-mers with no increasing pairs

In the histogram in Fig. 4bii, a higher number of peaks are observed on the decreasing side (specified by light blue) than on the increasing side (specified by dark blue), showing that many decreasing 15-mers were present. BLASTn analysis showed that these 15-mers were derived primarily from the beginning part of the 5’ UTR or the end part of the 3’ UTR. Many strains isolated in the early epidemic stage have been sequenced to near the start and end of the genome, but with the rapid pandemic spread, a large number of genomes for which the sequences in UTRs are undetermined have accumulated in the GISAID, and this had resulted in a time-series decrease for both end sequences. This decrease was due to artificial manipulation, and there was no relationship between heatmap patterns and clade territories (data not shown).

However, there was a peculiar group of 15-mers in the S gene, which showed a time-series decrease, but there was no pair showing an increase. In the heatmap, there was a characteristic blue pattern but not a red one (Fig. 4h); the small blue regions were scattered among multiple clade territories, showing that the genomes lacking the 15-mers of interest were scattered there. Since there was no paired red pattern in the heatmap and no pair increasing in the time-series analysis, the 15-mers decreasing in Fig. 4h may not be due to mutation. When the sequences located in the small blue areas were investigated, two consecutive bases in the S gene were registered as unknown bases (NN); CCUU (24980-24983) was registered as CNNU: N1 in Table1. When the total number of Ns in the genomes with the CNNU was examined, there was no tendency for the number to be higher than in other genomes. Therefore, this genomic site is considered to be difficult to sequence or inaccurate for sequencing; the CNNU sequence was observed in genomes of five different clades. Even in RNA viral genomes, modifications such as methylation are known to occur and their functions are drawing attention^20^, and some modifications may inhibit the proper progression of reverse transcriptase, resulting in sequencing difficulty. If the CNNU in the S gene is caused by RNA modification, the BLSOM will become a method for searching for such special genomic sites.

## Discussion

The analyses shown in Figs. 3 and 4 identified nine mutations, which were among the ten known mutations assigned by the phylogenetic clustering method based on sequence alignment. To explore the remaining one (denoted as M0 in Table 1), we analyzed time-series changes of fifteen 15-mers harboring the M0 mutation and found that their population frequency was 0.15 in February but only 0.09 in March and did not increase thereafter (data not shown). Because our analysis was based on 15-mers whose frequency increased/decreased to at least 0.1 in the March/June population, the M0 mutation was excluded from the analysis. To check this explanation, the increase/decrease level was reduced to 0.05, and the 2087 types of 15-mers obtained were used for additional BLSOM analysis (Supplementary Fig. S4). The reverse patterns of the red/blue heatmap and the time-series change were just as expected (data not shown), proving that the present AI method produce the result totally consistent with that of the phylogenetic clustering method. Notably, during AI assignment, the V clade separation was achieved even without the M0-related 15-mers, and this type of information should clarify the importance of individual mutations that contribute to each clade separation. The BLSOM of 2087 types of 15-mers with an increase/decrease of at least 0.05 should assign the mutations related to much more detailed subgroupings in each clade. Since the BLSOM has a powerful visualization ability, it can also provide information concerning genomic sites that are difficult to sequence (e.g., N1 in Table 1 and Fig. 4h) for various reasons, including possible RNA modifications.

Even during preparation of the present manuscript, many genome sequences of SARS-CoV-2 accumulated in the GISAID database. This appears to be a research difficulty but should provide a unique advantage. For the fast-evolving RNA virus, the near-future prediction and verification cycle can be realized, although the elementary process of molecular evolution is based on random mutation. This is because the time-series directional changes have been observed on a monthly basis most likely due to the viral adaptation for efficient growth in human cells. Near-future prediction and verification should be the most direct ways to test the reliability of the obtained results, models and ideas, providing a new paradigm for molecular evolutionary studies. Taking this view into consideration, we will proceed with the discussion.

In the present 15-mer analysis, we focused on the March/June population for the middle/final period and obtained 587 types of 15-mers that increased/decreased in population frequency by at least 0.1. When strains isolated later this year become available, our successive strategy will be to search for new 15-mers that increased/decreased their frequency by at least 0.1 (or 0.05) compared with that in December and construct a BLSOM of 15-mers including these newly obtained ones. A trial analysis of 3700 recently downloaded genomes isolated in August revealed only one new mutation that increased in frequency to 0.1 in the August population. This mutation had a frequency of 0.09 in June that increased to 0.12 in August; therefore, the corresponding 15-mers were included in the BLSOM presented in Supplementary Fig. S4. In the corresponding heatmap, red/blue inversion was observed within the GR territory, showing that it was an internal branch of GR. The mutation was an amino acid substitution: I120F in the gene encoding nsp2.

The phylogenetic method based on sequence alignment is a well-established and irreplaceable method for molecular evolutionary studies. The presently developed sequence alignment-free method is suitable for analyzing a massive amount of sequence data and can analyze over five million sequences simultaneously^8^; notably, this method is highly robust against sequencing errors, and therefore, no special pretreatment is required. Furthermore, the BLSOM is unsupervised AI that can be used without special models or presumptions, and has powerful visualization capabilities that enable efficient knowledge discovery from big data. Evolutionary studies have entered the era of big data, and the present method can complement phylogenetic methods based on sequence alignment, especially for a massive number of sequences.

## Methods

### Genome sequences

Genome sequences of human SARS-CoV-2 were downloaded from the GISAI database (https://www.gisaid.org/epiflu-applications/next-hcov-19-app/)^3^; sequences belonging to the complete genome, high-coverage and human categories were downloaded on July 20, 2020. We used all SARS-CoV-2 genome sequences (40,450) after removing their polyA-tails. Although a significant number of viral sequences contained many Ns (undetermined nucleotides), we did not conduct any special preprocessing because in big data analyses, such as word-count analyses, the effects of erroneous data appear to naturally decrease with an increase in dataset size. In the Discussion section, 3708 sequences isolated in August are additionally analyzed. The word-count program can be obtained from.

### BLSOM

The SOM developed by Kohonen et al. (1966)^21^ is an unsupervised neural network algorithm that implements characteristic nonlinear projection from the high-dimensional space of input data onto a two-dimensional array of weight vectors. We previously modified the conventional SOM for genome informatics to make the learning process and resulting map independent of data-input order and established a BLSOM^4^. The BLSOM for oligonucleotide composition was constructed as described previously^5^; initial vectorial data for the BLSOM were defined as the first and second components from PCA (principal component analysis). Because PCA can detect basic properties of genomic sequences, such as G+C%, global patterns of oligonucleotide BLSOMs, in which various learning parameters and the number of sequences per node are changed, resembled each other^5,7,16^. However, PCA requires a very long calculation time for k-mers of a large number of sequences (approximately 40,000 sequences in this case) when k>6. During machine learning, distance calculation is performed for sequence data and nodes (e.g., 1/50 of the number of sequences), and the calculation time is dramatically reduced. In other words, for 7-mer or longer oligonucleotides, the PCA process used only to determine the initial state requires a much longer computation time than machine learning, making efficient high-dimensional analysis difficult. To solve this problem of 7-mers in PCA, 1/100 of the sequences were randomly selected, and the PCA result obtained for these sequences was used to set the initial state. BLSOM programs were obtained from and http://bioinfo.ie.niigata-u.ac.jp/?BLSOMviewer#jc96a619.

## Acknowledgements

We gratefully acknowledge the authors submitting their sequences from GISAID’s Database. We gratefully acknowledge also the valuable comments of Dr. Yashushi Hiromi of National Institute of Genetics (Mishima). This work was supported by JSPS KAKENHI Grant Number 18K07151, by AMED under Grant Number JP20he0622033 and by COVID-19 Counterplan Research Project (supervised by Prof. Tatsumi Hirata, NIG) from the Research Organization of Information and Systems (ROIS).

## Author contributions

T.I. and T.A. designed this project and conducted genome analysis. K.W., Y.W. and Y.I. developed computer programs. T.I. wrote the manuscript. All authors participate in the discussion through the project.

## Additional information

Supplementary Figs. S1-S4.

